# Huntingtin expression influences spontaneous seizure disorder susceptibility in FVN/B mice

**DOI:** 10.1101/2022.10.18.512787

**Authors:** Jeremy M. Van Raamsdonk, Hilal Al-Shekeli, Laura Wagner, Tim W Bredy, Laura Chan, Jacqueline Pearson, Claudia Schwab, Zoe Murphy, Rebecca S. Devon, Ge Lu, Michael S. Kobor, Michael R. Hayden, Blair R. Leavitt

## Abstract

Huntington disease (HD) is an adult-onset neurodegenerative disorder that is caused by a trinucleotide CAG repeat expansion in the HTT gene that codes for the protein huntingtin (HTT or Htt in mice). HTT is a multi-functional, ubiquitously expressed protein that is essential for embryonic survival, normal neurodevelopment, and adult brain function. The ability of wild-type HTT to protect neurons against various forms of death raises the possibility that loss of normal HTT function may worsen disease progression in HD. Huntingtin-lowering therapeutics are being evaluated in clinical trials for HD, but concerns have been raised that decreasing wild-type HTT levels may have adverse effects. Here we show that Htt levels modulate the occurrence of an idiopathic seizure disorder that spontaneously occurs in FVB/N mice. These abnormal FVB/N mice demonstrate various cardinal features of mouse models of epilepsy including spontaneous seizures, astrocytosis, neuronal hypertrophy, upregulation of brain-derived neurotrophic factor (BDNF), and sudden seizure-related death. Interestingly, decreasing wild-type Htt levels increased the frequency of this disorder, while over-expression of HTT completely prevented it. Examination of the mechanism underlying huntingtin’s ability to modulate the frequency of this seizure disorder indicated that over-expression of full length HTT can promote neuronal survival following seizures. Overall, our results demonstrate a protective role for huntingtin in this form of epilepsy and provide a plausible explanation for the observation of seizures in the juvenile form of HD, Lopes-Maciel-Rodan syndrome, and Wolf-Hirschhorn syndrome. Adverse effects caused by altering huntingtin levels has ramifications related to Huntingtin-lowering therapies in development to treat HD.

## INTRODUCTION

Huntingtin (HTT) is a multi-functional protein, encoded by the huntingtin (*HTT*) gene located on chromosome 4p16.3. (1) HTT is ubiquitously expressed with the highest levels occurring in brain and testis. (2, 3) Individuals with an expanded (>35) CAG repeat in exon 1 of *HTT*, coding for a polymorphic polyglutamine tract, develop Huntington disease (HD) with an age of onset that is inversely related to the length of the CAG repeat expansion (OMIM #143100). (4) HD is an autosomal dominant neurodegenerative disorder that is characterized by motor dysfunction, cognitive impairment, and neuropsychiatric abnormalities (2). The early and relatively selective death of medium spiny striatal neurons in the brain is the predominant neuropathology reported in HD, manifesting before physical symptom onset. Individuals homozygous for an expanded CAG repeat in the HTT gene have more rapid disease progression than individuals heterozygous for the mutation. (5, 6)

Wild-type huntingtin function in mice is essential for embryonic survival (7–9), neurogenesis, (10–12) and for normal neuronal function during the postnatal period (13, 14). Decreasing Htt levels by 50% or more leads to neurological abnormalities in mice (8, 13, 15, 16) and makes mice more susceptible to toxic insults (17, 18). Huntingtin has been shown to have important functional roles in transcription, intracellular transport, and neuroprotection (19). Wild-type Htt expression has been shown to influence the pathogenesis of HD mouse models (20–22). Multiple huntingtin-lowering therapeutic agents targeting both wild-type and mutant HTT have made it to clinical trials. (23) It is currently uncertain whether wild-type HTT reduction may alter the potential benefit of mutant huntingtin lowering or even have adverse effects in the adult CNS.

Individuals with > 60 CAG repeats in *HTT* usually develop a juvenile-onset form of HD, with onset typically by the age of 20. Juvenile-onset HD is characterized by a more severe disease progression and a different variation of symptoms that are not present in the adult form of HD, including an increased incidence of epilepsy. (24, 25) In fact, the longer the CAG repeat expansion and the younger the age of onset, the greater the likelihood that a person with HD patient would exhibit seizures. (25, 26) As in the human disease, seizures have been observed in several mouse models of HD with very large CAG repeat expansions. The R6/2 mouse model, which expresses an N-terminal fragment of mutant Htt (27), has been shown to have increased susceptibility to both chemically-induced and audiogenic seizures (28, 29). As R6/2 mice also have decreased levels of full-length wild-type Htt (30, 31), it is possible that decreased neuroprotection by wild-type Htt may contribute to the development of epilepsy in this model. In support of this hypothesis, it has been shown that epilepsy is a common feature of Lopes-Maciel-Rodan syndrome (LOMARS) (OMIM #617435) and Wolf-Hirschhorn syndrome (OMIM #194190). LOMARS patients have a RETT syndrome-like phenotype caused by compound heterozygous mutations in the *HTT* gene. (32, 33) The neurodevelopmental phenotype in LOMARS is thought to be due to the co-existence of null and hypomorphic *HTT* alelles. (34) Wolf-Hirschhorn syndrome is a congenital malformation syndrome, which results from a hemizygous deletion on chromosome 4p16.3 that includes the *HTT* gene. (35) Furthermore, mouse models of Wolf-Hirschhorn syndrome, with decreased Htt levels, also exhibit increased susceptibility to seizures (36).

To investigate the role of Htt in the development of epilepsy in mice, we examined the effect of altering huntingtin levels on a novel, idiopathic seizure disorder that occurs in FVB/N mice. We provide a thorough characterization of the seizure disorder and demonstrate that decreasing expression levels of Htt increases frequency of this disorder. Over-expression of HTT decreases the frequency of the seizure disorder and the amount of seizure-induced neurodegeneration. These results suggest that huntingtin protects against seizures.

## RESULTS

### A novel seizure disorder in FVB/N mice

To evaluate a potential role of huntingtin in preventing the development of epilepsy, we utilized a novel idiopathic seizure disorder model identified in our laboratory. Among FVB/N mice, we observed that a proportion of wild-type (WT) mice exhibit home cage immobility coupled with abnormal aversion to handling that resembles the post-ictal state of mice with chemically-induced seizures (**Movies 1-3**). These mice are genetically wild-type (100%congenic to littermates). To determine whether the post-ictal appearance of these abnormal WT FVN/N mice resulted from spontaneous seizures, we monitored abnormal and normal WT FVB/N mice for seizure activity for a period of two hours. Mice were categorized as abnormal FVB/N mice based on home cage immobility and aggressive response to handling with post-mortem confirmation by brain pathology (see below). During the two-hour monitoring period we found that 11 of 14 abnormal WT FVB/N mice exhibited spontaneous seizures with a total of 22 seizures (see **Movie 4** for an example of a typical seizure observed). In contrast, no seizures were observed in normal WT FVB/N mice.

As mouse models of epilepsy have been shown to exhibit increased sensitivity to induced seizures (37), we examined the susceptibility of these abnormal WT FVB/N mice to audiogenic seizures and chemical induction of seizure with PTZ. After exposure to an audiogenic stimulus, 60% of the abnormal WT FVB/N mice exhibited a “pop-corn” seizure followed by immediate death (**Movie 5**). In contrast, none of their normal WT FVB/N littermates seized or reacted in any way to the audiogenic stimulus (**Movie 6**). This phenomenon of sudden unexpected death in epilepsy (SUDEP) occurs in multiple forms of human epilepsy but the underlying mechanisms are not currently known (38, 39). Chemical induction of seizures with pilocarpine also led to SUDEP in 66% of the abnormal WT FVB/N mice, with no effect on normal WT FVB/N littermate controls. Based on this phenotype, we have named the abnormal WT FVB/N mice **FSDS mice** (FVB/N Seizure Disorder with SUDEP).

The average age of onset for the FSDS phenotype was 6.5 months and was characterized by altered home cage activity, aversion to handling and refusal to participate in routine behavioural tests such as rotarod or beam crossing. FSDS mice also exhibit a bi-phasic pattern of weight change consisting of a mild increase in body weight around the time of onset that is followed by a dramatic decrease in weight of up to 15 grams. Finally, FSDS mice exhibit severely abnormal activity patterns in an automated open field activity test with periods of both hyperactivity and hypoactivity (**Fig. 1A,B**).

**Figure 1.**
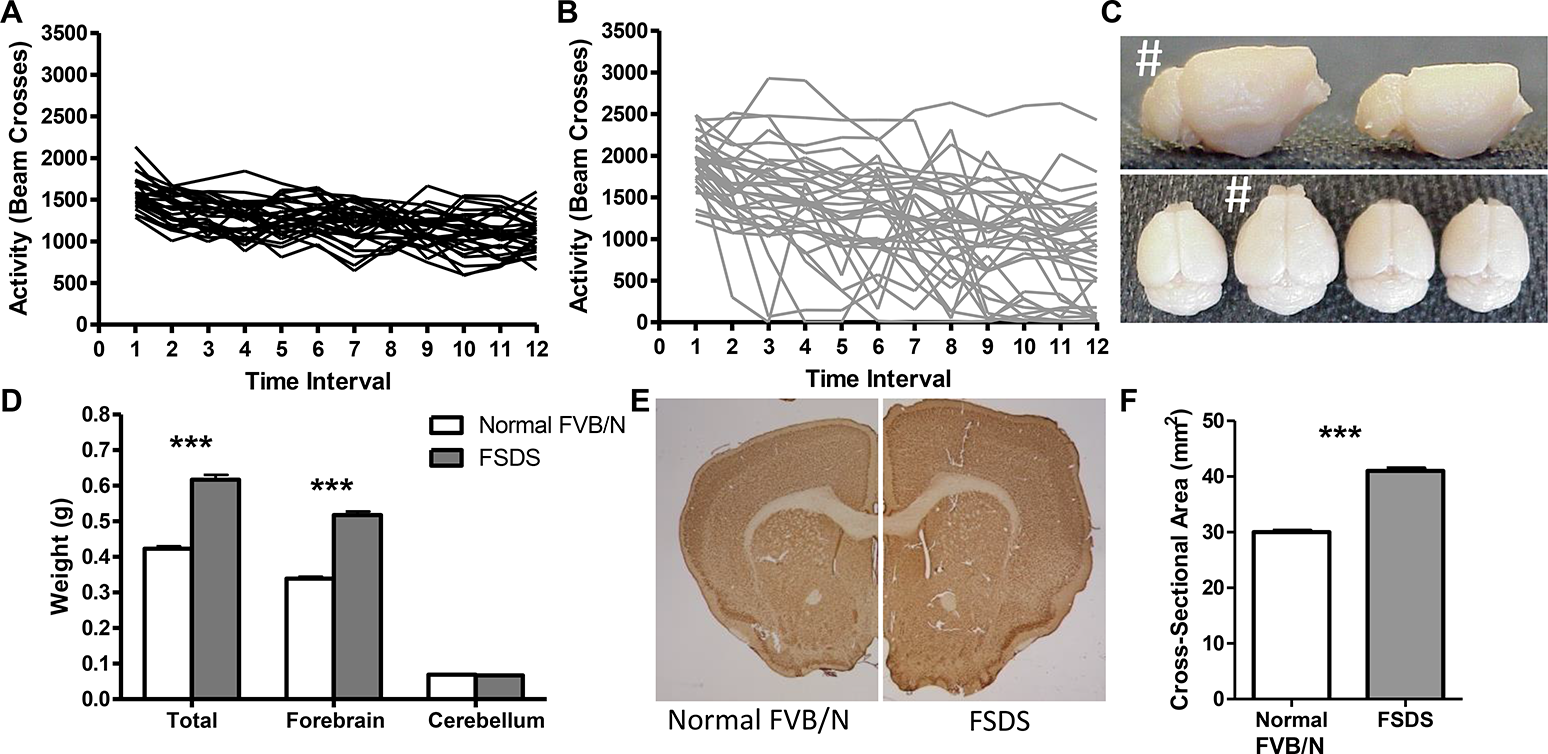
FVB/N Seizure Disorder with SUDEP (FSDS) mice exhibit abnormal activity and megacephaly. A, Normal FVB/N mice (N = 28) exhibited a uniform pattern of activity where activity declines over the one hour open field trial (each line represents one mouse). B, In contrast, FSDS mice (N = 31) exhibited multiple periods of both hypoactivity and hyperactivity. C, The brains of FSDS mice (shown with # sign) were much larger than the brains of normal FVB/N mice. D, FSDS mice exhibited a 46% increase in brain weight resulting only from an increase in the forebrain, while the cerebellum is spared (N = 12 per group). E-F, The brains of FSDS mice also exhibited a significant increase in brain cross-sectional area compared to normal FVB/N mice (N = 6 per group). Error bars indicate SEM. *** p<0.001.

### FSDS phenotype does not result from a spontaneous mutation

To determine whether the FSDS phenotype resulted from a spontaneous mutation in the FVB/N strain, we examined whether the offspring of FSDS mice are more likely to develop the FSDS phenotype than the offspring of normal WT FVB/N mice. We found that the offspring of FSDS showed no enrichment for the FSDS phenotype, suggesting that the phenotype does not result from a dominant mutation. In crossing two FSDS mice together, we were unable to establish an FSDS line, suggesting that the phenotype does not result from a recessive mutation (i.e. does not breed true). Conversely, we attempted to breed out the FSDS phenotype by only breeding offspring of normal WT FVB/N mice. Even after multiple generations, we were unable to eliminate the FSDS phenotype. Combined, these results suggest that the FSDS phenotype does not result from a spontaneous mutation. Further support for this conclusion comes from the fact that we have observed this phenotype in two different animal facilities where the WT FVB/N colonies are completely independent. The observation of the FSDS phenotype at two independent animal facilities also suggests that the phenotype can arise in multiple environments. Furthermore, we have observed this phenotype in FVB/N mice obtained from both Jackson Laboratories and Charles River. As we have never observed this phenotype in other strain backgrounds, this suggests that the FVB/N strain has a genetic susceptibility to the development of the FSDS phenotype.

### FSDS mice exhibit megacephaly, electrographic seizures, astrocytosis and neuronal hypertrophy

To determine whether the epileptic phenotype in FSDS mice resulted from changes in brain morphology, we compared the brains of FSDS mice and normal FVB/N littermates. It was immediately apparent that the brains of the FSDS mice were abnormal as they were found to be grossly enlarged (**Fig. 1C**). Quantification revealed that the brains weighed almost 50% more than normal age-matched FVB/N control mice, with no enlargement observed in the cerebellum (**Fig. 1D**; whole brain - normal FVB/N: 423 ± 6 mg, FSDS: 617 ± 13 mg, p < 0.001; cerebellum - normal FVB/N: 69 ± 1 mg, FSDS: 67 ± 1 mg, p = 0.28). In addition, FSDS brain sections exhibited a 37% increase in cross-sectional area compared to normal littermate controls (**Fig. 1E,F**; normal FVB/N: 30.0 ± 0.2 mm^2^, FSDS: 41.0 ± 0.4 mm^2^, p < 0.001).

To determine whether the seizure activity that was observed behaviourally is associated with underlying electrophysiological abnormalities, a total of 103 and 130 hours of multi-channel EEG recordings were obtained form the FSDS and normal WT FVB/N mice, respectively. Analysis of EEG data from FSDS mice revealed the presence of recurrent epileptiform discharges. The discharges, which were synchronous in all EEG channels, correlated with abrupt and intense movement changes along the three accelerometer axes, indicative of a convulsive seizure. Typical EEGs during the ictal discharges are shown in **Fig. 2**. The seizure initiated with a burst of 400-μV spikes that progressively become faster reaching a frequency of 18 Hz (**Fig. 2C ii**). The spikes amplitude also escalated with time until reaching about 990 μV at the peak of the episode (**Fig. 2C iii**). Post- and inter-ictal spikes were also frequently observed. Duration of the ictal phase ranged from 13 to 42 seconds, while the full electrographic phase lasted for up to 64 seconds. Ictal discharges occurred at a rate of 22 times per 29 hours of continuous EEG recording in the first mouse, compared to 18 times per 43 hours in the second mouse. Seizures seem to occur in clusters where the shortest time between two successive seizures was 157 seconds and the longest seizure-free interval was 12.5 hours. No similar epileptiform discharges were noted in the EEG recordings from normal WT mice.

**Figure 2.**
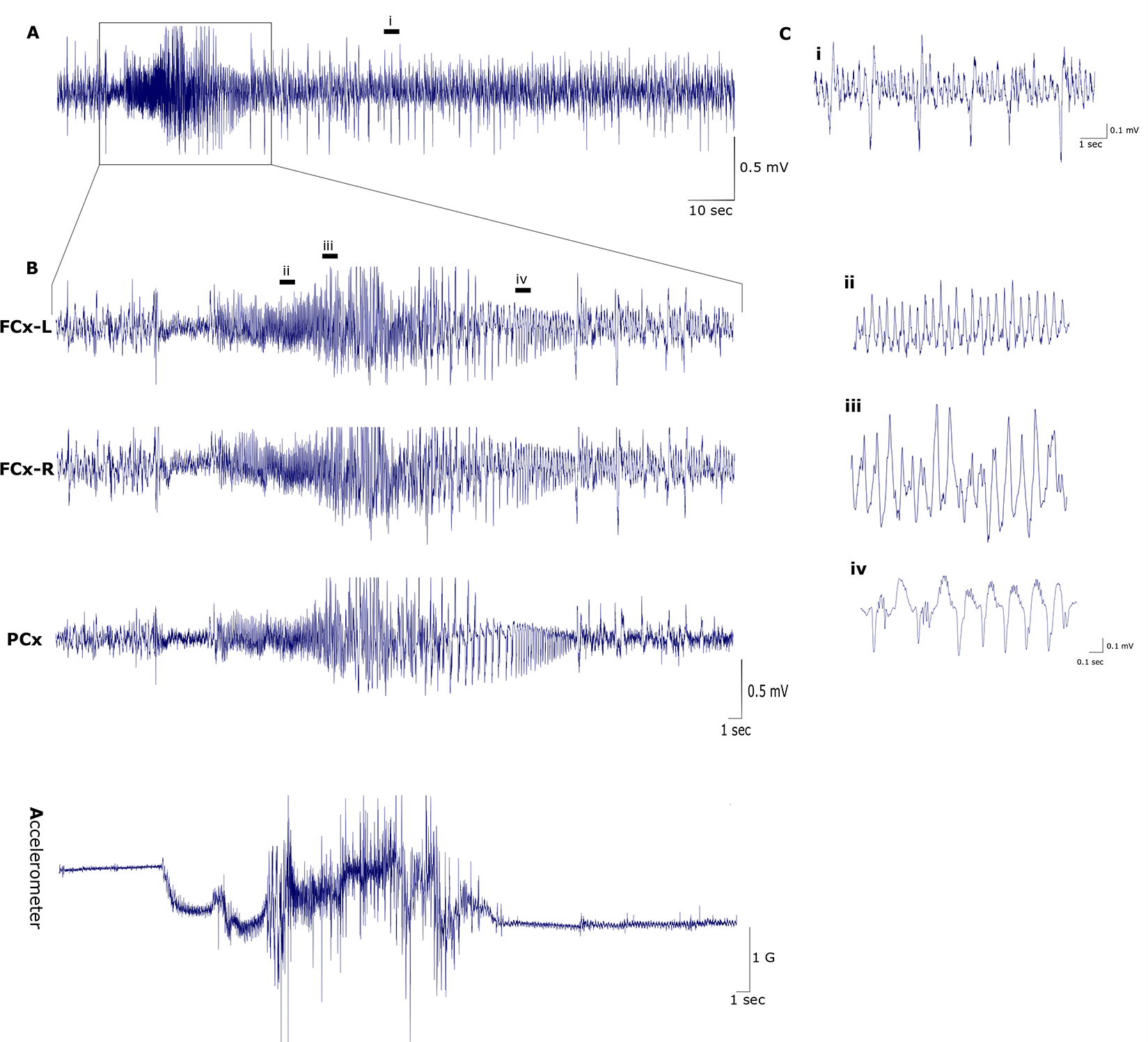
Representative EEG from FSDS mice showing epileptiform discharges. A, condensed view of 145-seconds epoch showing a burst of high-amplitude discharges. B, expanded views of the burst showing synchronous ictal discharges from the 3 EEG channels along with concurrent trace from accelerometer x-axis. The onset of the discharges coincides with changes in the accelerometer that becomes more intense with time and then tapers down and flattens towards the end of the episode. C, further expanded views showing waveform morphology of post-ictal spikes (i) and in 8-seconds samples taken form the beginning (ii), middle (iii) and end (iv) of the epoch. Abbreviations, FCx-L: left frontal cortex, FCx-R: right frontal cortex, PCx: parietal cortex, sec: seconds, mV: millivolts, G: acceleration of Earth’s gravity (~9.8 m/s).

Having shown gross brain abnormalities and electrophysiological differences in FSDS mice, we sought to determine whether these mice exhibited cellular neuropathology consistent with seizure disorders. Accordingly, we examined FSDS mouse brains for astrocytosis, which has been observed in mouse models of epilepsy (40). For comparison, we also examined the brains of mice from two chemically induced models of seizures: pilocarpine-induced SE and PTZ. Pilocarpine is a non-subtype-specific partial muscarinic agonist that has been used to model temporal lobe epileptic seizures in humans (41), while PTZ is a GABA receptor antagonist and is used as a model of generalized epilepsy (42).

As in the pilocarpine and PTZ models (not shown), we found dramatic astrocytosis in the brains of FSDS mice, which was not observed in normal FVB/N controls (**Fig. 3A**). Next, we sought to determine whether the astrocytosis exhibited regional specificity. We found that astrocytosis was limited to regions of the brain comprising the limbic system (hippocampus, amygdala and piriform cortex), which is typically affected in temporal lobe epilepsy while other regions, such as the striatum, were unaffected. As with patients with temporal lobe epilepsy (43), we also observed selective neuronal hypertrophy in affected regions: the piriform cortex and amygdala but again not the striatum (**Fig. 3B,C**; piriform cortex - normal FVB/N: 152 ± 3 μm^2^, FSDS: 187 ± 6 μm^2^, p = 0.01; amygdala - normal FVB/N: 145 ± 2 μm^2^, FSDS: 207 ± 10 μm^2^, p = 0.02; striatum - normal FVB/N: 64 ± 1 μm^2^, FSDS: 66 ± 1 μm^2^, p = 0.2).

**Figure 3.**
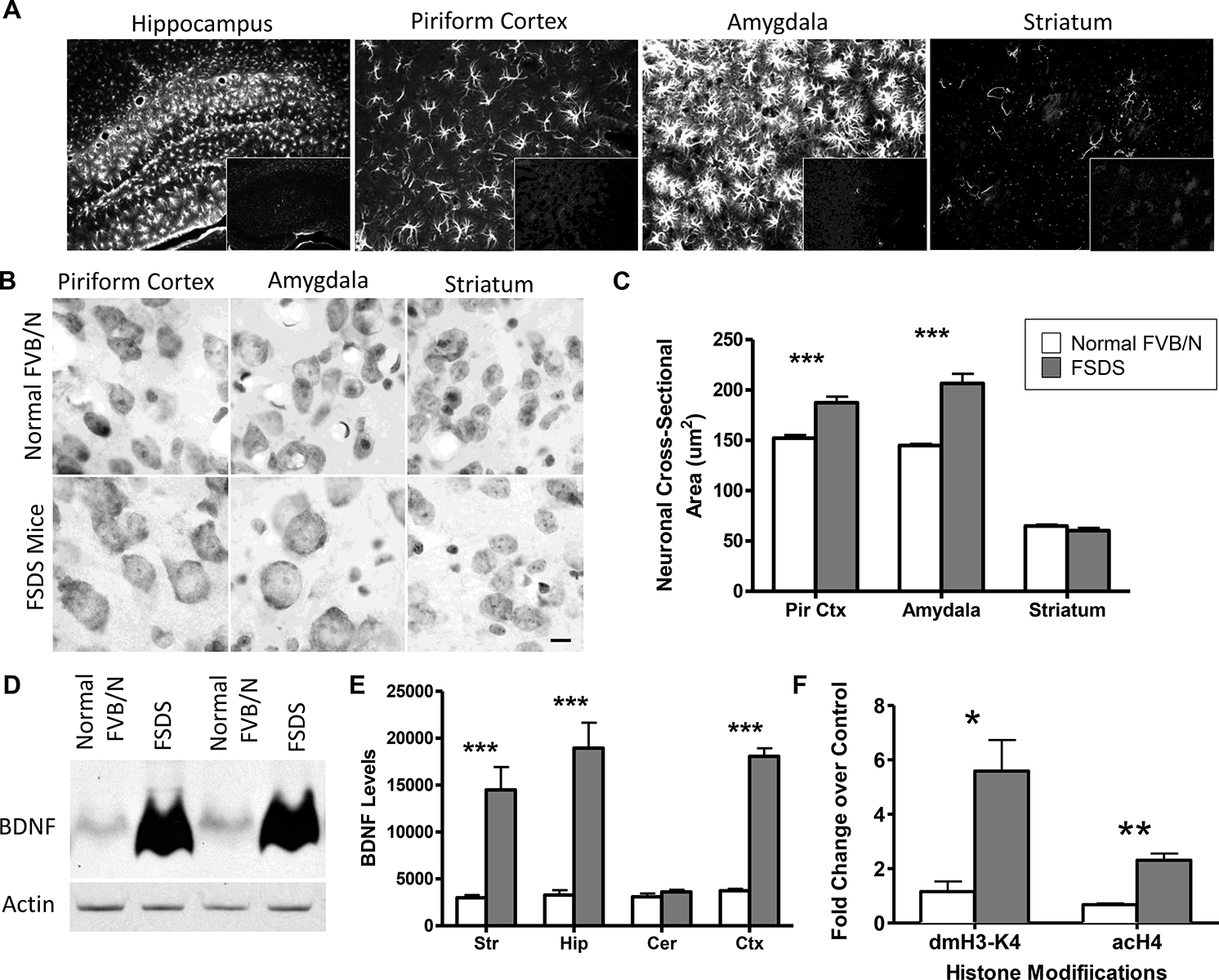
Brains from FSDS mice exhibit neuropathology characteristic of mouse models of epilepsy. A, Immunostaining with a Cy3-labelled anti-GFAP antibody reveals that FSDS mice exhibit astrocytosis selectively in hippocampus, piriform cortex and amygdala (large frame) but not the striatum, while minimal glial fibrillary acidic protein staining is observed in normal FVB/N mice (small inset frame). B-C, FSDS mice also showed neuronal hypertrophy selectively in piriform cortex and amygdala but not the striatum (N = 3 per group). D, At the molecular level, FSDS mice showed a dramatic increase in brain-derived neurotrophic factor (BDNF) expression by Western blotting. E, The increase in BDNF levels was only observed in forebrain regions - not the cerebellum (N = 3 per group). F, Epigenetic changes at BDNF promoter 2 (dimethylation of H3-K4, acetylation of H4) contribute to the increased BDNF expression in FSDS mice (N = 3 per group). Error bars indicate SEM. * p<0.05, ** p<0.01, *** p<0.001.

### FSDS mice exhibit marked upregulation of BDNF resulting from epigenetic changes at the BDNF promoter

At the molecular level, seizure activity has been shown to result in increased levels of BDNF (44, 45). In addition, increased levels of BDNF have been shown to promote seizure activity (46–48). To determine whether the FSDS phenotype is associated with increased expression of BDNF, we measured the levels of BDNF protein in the brains of 12 month old FSDS mice and found a dramatic increase in BDNF expression similar or greater to that observed in two chemically induced models of epilepsy (**Fig. 3D**; normal FVB/N: 3226 ± 343 arbitrary units; FSDS: 13677 ± 417 arbitrary units, p < 0.001).

To determine whether the increase in BDNF expression occurs in specific regions of the brain, we micro-dissected brains from FSDS and normal FVB/N mice and examined brain region specific BDNF protein levels by Western blotting. While the levels of BDNF were similar in the striatum, hippocampus, cortex and cerebellum in normal FVB/N mice, the increased BDNF expression in FSDS mice was limited to the striatum, hippocampus and cortex (**Fig. 3E**). Examination of *Bdnf* mRNA levels indicated that the increase in BDNF protein levels resulted from increased transcription of *Bdnf* mRNA (not shown).

Next, we sought to determine whether the increase in BDNF expression in FSDS mice resulted from epigenetic changes at the BDNF promoter as the acetylation of histone H4 (H4 acetylation) has previously been observed at the BDNF promoter in chemical and electroconvulsive seizure models (49, 50). We also measured dimethylation of lysine 4 on histone H3 (H3-K4 dimethylation) as this modification has been associated with transcriptional activation (51). Examination of BDNF promoter 2 revealed a 5-fold increase in H3-K4 dimethylation and 3-fold increase in H4 acetylation in FSDS mice compared to normal FVB/N mice (**Fig. 3F**). As both of these modifications have been associated with transcriptional activation, this suggests that epigenetic changes at the BDNF promoter contribute to the initiation and or maintenance of increased BDNF expression in FSDS mice.

### Full-length wild-type huntingtin levels modulate the frequency of FSDS phenotype

Having shown that FSDS mice exhibit characteristic features of mouse models of epilepsy, we sought to determine whether wild-type Htt function would protect against the development of epilepsy in these mice. We have previously shown that over-expression of wild-type HTT reduces seizure-induced neurodegeneration caused by the delivery of either the glutamate receptor agonist kainic acid or the NMDA-specific glutamate receptor agonist quinolinic acid, in the hippocampus or striatum respectively (30, 52).

To assess the effect of Htt levels on the development of the FSDS phenotype, we compared the frequency of the FSDS phenotype between mice heterozygous for the targeted inactivation of the mouse *Htt* gene (*Htt*+/− mice) and their WT littermates. While the *Htt*+/− mice were originally generated on a S129/C57BL/6 strain background (8), they were subsequently backcrossed onto the FVB/N strain background for more than 10 generations. Mice were categorized as FSDS mice based on home cage immobility, aggressive response to handling and megencephaly post-mortem (**Fig. 4A**). In each case, we observed a perfect correlation between the behavioural symptoms and brain pathology. In comparing the frequency of the FSDS phenotype between *Htt*+/− mice and their WT littermates, we found that 24% of the WT littermates developed the FSDS phenotype by 12 months of age (**Fig. 4B**, **Table 1**). In contrast, *Htt*+/− mice, which express Htt at approximately 50% of wild-type levels, developed the FSDS phenotype at a frequency of 71%, despite being born from the same parents and being genetically identical except at the *Htt* gene locus (**Fig. 4B**; χ^2^ = 10.0, p = 0.0015).

**Table 1.** Increased huntingtin expression is associated with decreased frequency of FSDS mice. Mice were followed for 1 year. FSDS phenotype was determined by observation of abnormal home cage behaviour that included inactivity and aggressive response to handling. FSDS phenotype was confirmed by observation of megencephaly post-mortem.

| Genotype | Copies of Mouse *Htt* Gene | Copies of Full-Length *HTT* Transgene | Copies of full length *HTT* gene | Total Mice Examined | Number of FSDS Mice | Frequency of FSDS Mice | Difference from WT |
| --- | --- | --- | --- | --- | --- | --- | --- |
| *Htt +/-* | 1 | 0 | 1 | 14 | 10 | 71% | $\chi^2 = 10.0$, $p = 0.0015$ |
| WT | 2 | 0 | 2 | 95 | 27 | 28% | N/A |
| ShortStop | 2 | 0 | 2 | 14 | 5 | 36% | $\chi^2 = 0.3$, $p = 0.6$ NS |
| YAC18 | 2 | $\geq 1$ | $\geq 3$ | 38 | 0 | 0% | $\chi^2 = 13.6$, $p = 0.0002$ |
| YAC128 | 2 | $\geq 1$ | $\geq 3$ | 56 | 0 | 0% | $\chi^2 = 19.4$, $p < 0.0001$ |

**Figure 4.**
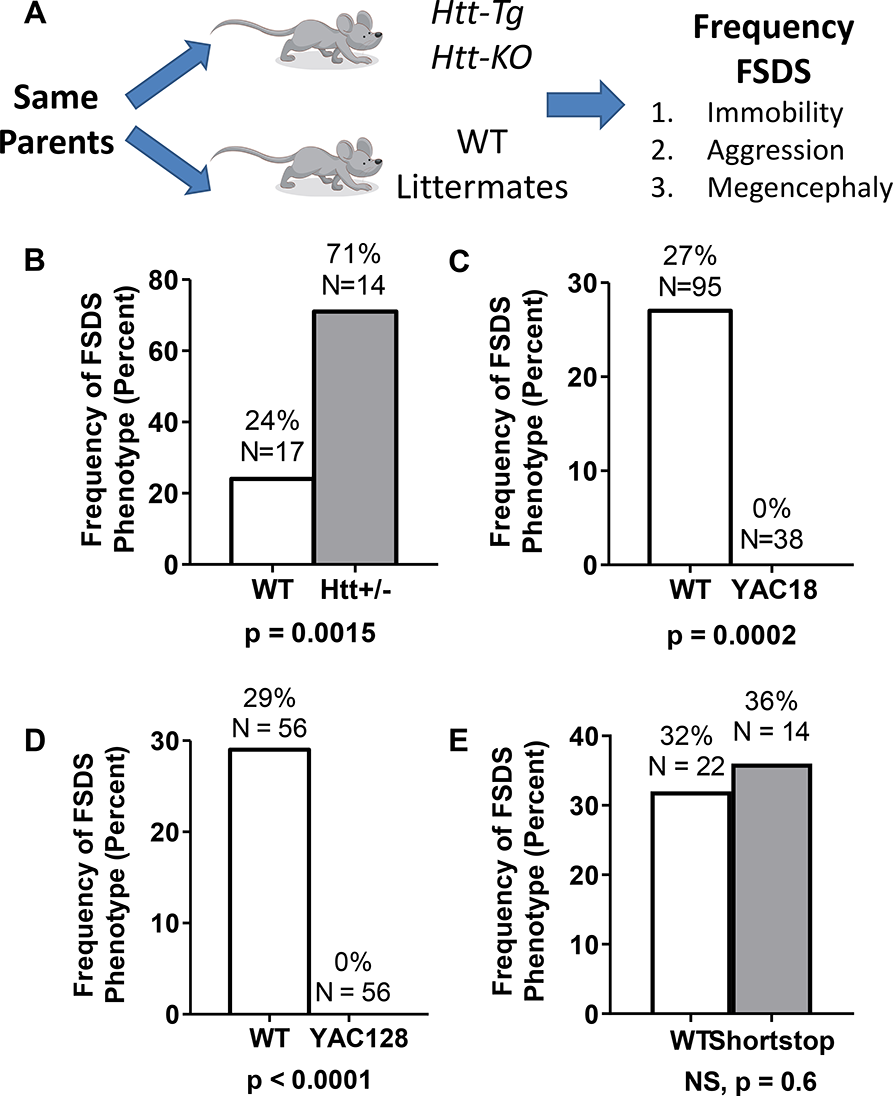
Huntingtin expression modulates the frequency of the FSDS epileptic phenotype. A, To examine the effect of Htt expression on the development of the FSDS phenotype, we compared the FSDS frequency between mice with altered levels of Htt and their wild-type littermates. FSDS mice were identified based on home cage immobility, an aggressive response to handling and megacephaly post-mortem. B, A 50% reduction in huntingtin (Htt) levels in *Htt*+/− mice more than doubled the frequency of the FSDS phenotype. C, Conversely, increasing the expression of full length wild-type HTT completely eliminated the occurrence of the FSDS phenotype in YAC18 mice. D, Similarly, increased expression of full-length mutant HTT completely prevented the development of the FSDS phenotype in YAC128 mice. E, In contrast, expression of an N-terminal fragment of mutant HTT in shortstop mice had no effect on the frequency of the FSDS phenotype. The data and statistical analyses for panel A are presented in Table 1. Note that in panels B, D and E littermate controls were used for WT animals. Panel C uses a composite of WT animals from B, D, and E. Htt-Tg = transgenic mice expressing increased levels of full length or mutant HTT. Htt-KO = heterozygote Htt knockout mice.

Since normal levels of wild-type Htt protect against the development of the FSDS phenotype in WT mice, we next sought to determine whether HTT over-expression would provide further protection. For this purpose, we examined the frequency of the FSDS phenotype in YAC18 mice (53), which over-express human HTT with 18 CAG repeats at approximately 2–3 times endogenous levels on an FVB/N strain background. We found that over-expression of wild-type HTT completely prevented the occurrence of the FSDS phenotype (**Fig. 4C**; **Table 1**; χ^2^ = 13.6, p = 0.0002). This indicates that increased levels of wild-type HTT protect against the development of the FSDS epileptic phenotype.

### Full-length mutant huntingtin, but not mutant huntingtin fragments, prevents the development of the FSDS phenotype

As we have previously shown that mutant HTT can replace the function of wild-type Htt during development (20) and in influencing body weight (54), we next sought to determine whether the expression of full length mutant HTT could prevent the development of the FSDS phenotype. For this purpose, we used YAC128 mice, which over-express full-length mutant human HTT with 128 CAG repeats at approximately 3/4 of wild-type levels (total full-length huntingtin levels are approximately 175% of wild-type levels) on an FVB/N strain background (55). We found that YAC128 mice never develop the FSDS phenotype while their WT littermates developed the FSDS phenotype at a rate of 29% (**Fig. 4D**; **Table 1**; χ^2^ = 19.4, p < 0.0001).

To determine whether the ability of HTT to modulate the frequency of the FSDS phenotype was a function of the full-length huntingtin protein, we examined the development of the FSDS phenotype in shortstop mice. Shortstop mice over-express only exons 1 and 2 of mutant HTT with 128 CAG repeats from the same yeast artificial chromosome (YAC) that was used to generate YAC128 mice and were generated on an FVB/N strain background (56). These mice have reported similar full length Htt levels as wild-type mice, while there is increased full length huntingtin protein in the YAC18 and YAC128 models. (53, 55, 56) In contrast to what was observed in YAC128 mice, we found that shortstop mice develop the FSDS phenotype at a frequency equivalent to their WT littermates (36% vs. 32%; **Fig. 4E**; **Table 1**; χ^2^ = 0.1, p > 0.05). This suggests that protection against the FSDS phenotype is not mediated by regions exon 1 or 2 in *HTT*. In addition, this excludes the possibility that the protective effect observed in YAC18 and YAC128 mice results from the over-expression of any transgene from a YAC and instead indicates that over-expression of both mutant and wild-type HTT specifically prevents the development of the FSDS phenotype. Additionally, it suggests that mutant huntingtin retains some of huntingtin’s normal protective function.

Next, we sought to determine whether we could increase the frequency of the FSDS phenotype using treatments that have previously been shown to induce seizure activity. Unlike decreasing Htt levels, we were unable to increase the frequency of the FSDS phenotype with handling-stress (57), induction of inflammation (58), or repeated exposure to auditory stimuli (59). There was also no effect of gender on the FSDS phenotype as both males and females WT mice developed this phenotype equally (males: 14/32, 30% FSDS; females: 13/36, 27% FSDS; p = 0.5 NS).

### Huntingtin expression levels are normal in FSDS mice

Based on the ability of huntingtin to modulate the frequency of the FSDS phenotype, we examined the levels of Htt protein in FSDS and normal FVB/N mice to determine if Htt levels might be decreased in FSDS mice. We found that Htt levels were not altered in the brains of FSDS mice compared to normal FVB/N littermates (**Fig. 5**). This indicates that while the levels of full-length Htt clearly modulate the frequency of the FSDS phenotype (**Fig. 4A-E**, **Table 1**), that the FSDS phenotype does not result from differences in Htt expression. Thus, decreased levels of Htt in wild-type FVB/N mice do not cause the FSDS phenotype, rather the expression of Htt protects against the development of this phenotype.

**Figure 5.**
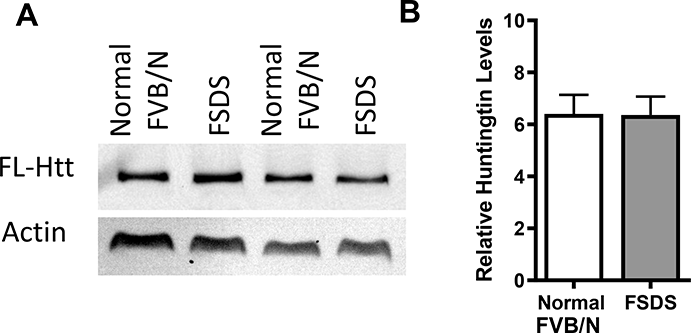
Huntingtin levels are unaltered in FSDS mice. Since the levels of full-length Htt clearly modulate the frequency of development of the FSDS phenotype, we sought to determine whether FSDS mice have reduced levels of Htt compared to normal FVB/N mice. A-B, Western blotting and quantification of Htt levels revealed no difference between normal FVB/N mice (N=9) and FSDS mice(N=7). Htt levels were normalized to actin. This indicates that while decreased levels of Htt increase the frequency of the FSDS phenotype, mice that develop the FSDS phenotype do not have decreased levels of Htt.

### Huntingtin limits seizure induced neurodegeneration but does not reduce frequency or severity of seizures

Given the clear impact of full-length huntingtin levels on the development of the FSDS epilepsy phenotype, we next examined possible mechanisms by which HTT provides protection. In order to determine whether huntingtin’s protective effect might result from a decrease in seizure susceptibility or a decrease in damage resulting from seizure, we utilized the PTZ kindling model of epilepsy to examine susceptibility, and pilocarpine-induced SE to examine neuronal damage caused by seizure. A comparison between mice over-expressing wild-type HTT (YAC18 mice, N = 19) and their WT littermate controls (N = 17) after repeated injections of PTZ revealed no difference in the number or severity of seizures induced by kindling (**Fig. 6A,B**). Similarly, injection of pilocarpine resulted in SE in equal proportions in WT and YAC18 mice (WT: 6 of 12, YAC18: 4 of 9, p = 0.15). The fact that HTT does not impact the number or severity of seizures in either chemically induced seizure model suggests that over-expression of HTT does not protect against the FSDS phenotype by preventing seizures or limiting seizure severity.

**Figure 6.**
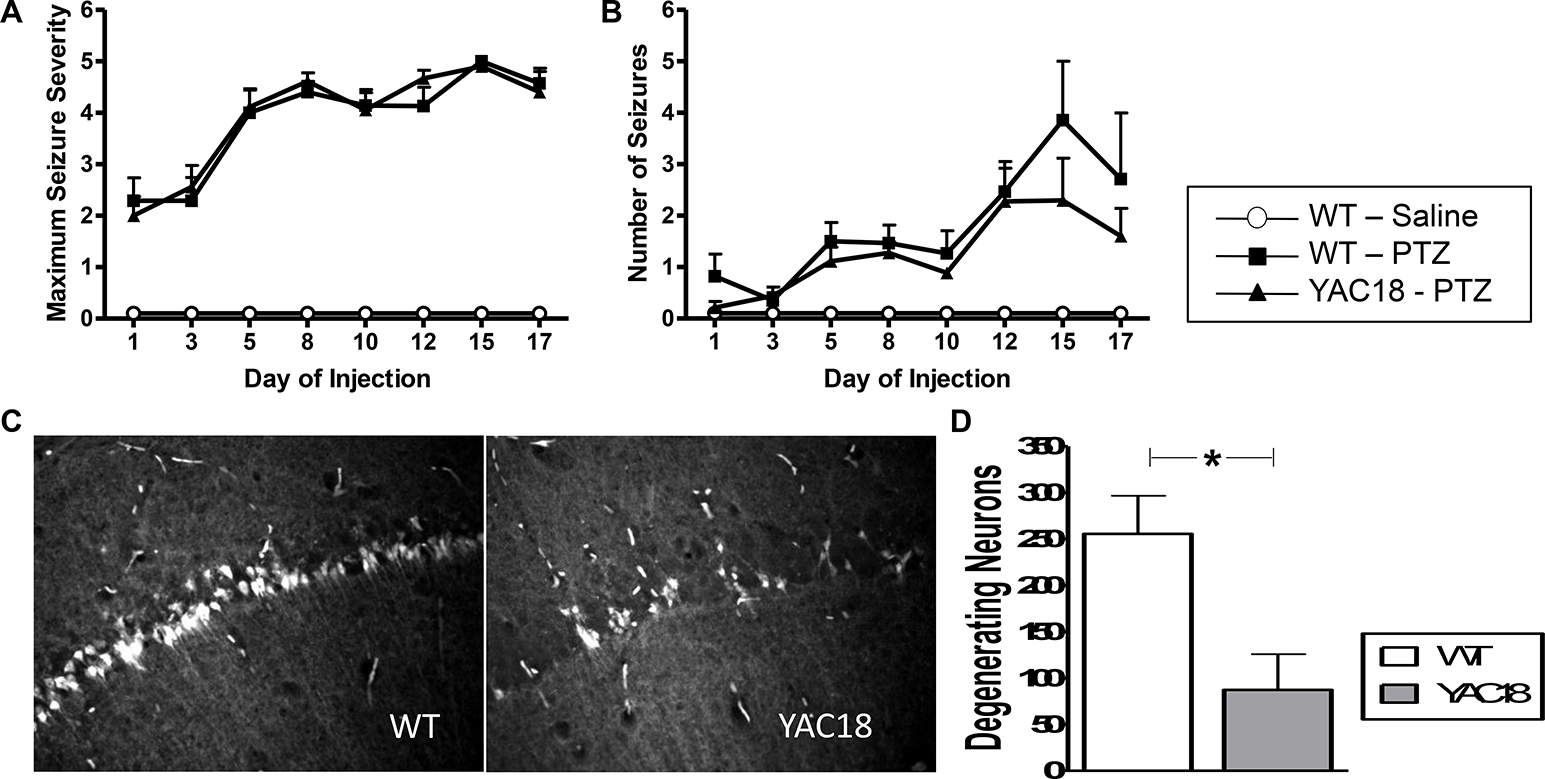
Over-expression of huntingtin reduces seizure-induced neurodegeneration. To assess seizure susceptibility, WT mice (N =17) and YAC18 mice (N=19) were given repeated injections of pentylenetetrazole (PTZ). WT mice (N=7) were also injected with saline as a control. A-B, Over-expression of huntingtin (HTT) had no effect on either the maximum seizure severity or the total number of seizures. To assess seizure-induced neurodegeneration, WT mice (N=12) and YAC18 mice (N = 9) were injected with pilocarpine. Sections containing the hippocampus were stained for degenerating neurons using fluorojade. C-D, Among those mice that developed status epilepticus (6 WT, 4 YAC18), mice over-expressing HTT showed significantly decreased numbers of degenerating neurons. Error bars indicate SEM. * p<0.05.

Finally, we examined the effect of HTT on seizure induced neurodegeneration. Of the mice that had undergone SE in response to pilocarpine, we stained a series of coronal sections throughout the hippocampus with fluorojade to detect degenerating neurons (**Fig. 6C**). While over-expression of HTT had no effect on seizure number or severity, mice over-expressing wild-type HTT exhibited significantly less neuronal damage following SE than WT controls (**Fig. 6D**; WT: 255 ± 41 degenerating neurons/section, YAC18: 87 ± 38 degenerating neurons/section, p = 0.02). This suggests that over-expression of HTT may protect against the FSDS phenotype by reducing seizure induced neurodegeneration.

## DISCUSSION

The development of epileptic seizures is a common feature of the juvenile form of HD and the likelihood of developing seizures increases with the number of CAG repeats present in the mutant *HTT* gene (25, 26). An increased susceptibility to seizures has also been observed in the R6/2 mouse model of HD (28, 29), which has decreased levels of wild-type Htt and a polyglutamine expansion that would fall within the juvenile range (27). Based on our previous work that demonstrates that wild-type HTT is neuroprotective (20, 21, 30, 52), and the fact that full length Htt levels are decreased in R6/2 mice (30, 31), we hypothesized that the levels of full length Htt could modulate susceptibility to the development of seizures. To study the effect of Htt expression on the development of seizures, we first characterized a novel mouse model of idiopathic epilepsy, which we have called <u>F</u>VB/N <u>S</u>eizure <u>D</u>isorder with <u>S</u>UDEP or FSDS. We demonstrate that the expression levels of wild-type or mutant huntingtin modulate the frequency of the FSDS phenotype and over-expression of HTT reduces seizure-induced neurodegeneration.

FSDS mice exhibit characteristic features of mouse models of epilepsy

FSDS mice are abnormal FVB/N mice that are genetically wild-type and occur at a frequency of ~30%. FSDS mice are identified by home cage immobility and aversion to handling, followed by confirmation of megacephaly at death. We observed a perfect correlation between behavioural abnormalities and brain enlargement. FSDS mice exhibit several features that have been observed in mouse models of epilepsy including spontaneous seizures, increased susceptibility to induced seizures (37), astrocytosis (40), neuronal hypertrophy (43) and upregulation of BDNF expression (44, 45).

One of the most striking features of FSDS mice is their dramatic increase in brain size. The megacephaly mouse (*mceph/mceph*) has an 11 base pair deletion in a voltage gated potassium channel (*KCNA1*) that results in a 25% increase in brain size (60). These mutants arose from a spontaneous mutation on the BALB/c strain background and, as with the FSDS mice, exhibit spontaneous seizures, increased levels of *Bdnf* mRNA and astrocytosis. In contrast to FSDS, the seizure phenotype in *mceph* mice is mild (less seizures, mild astrocytosis) and the age of onset is much earlier, with abnormalities apparent at 3 weeks of age (60). The fact that treatment of these mice with carbamazepine limits brain overgrowth without reducing seizures indicates that the increase in brain size is not the cause of the epileptic phenotype (61). The conclusion that an increase in brain size alone is insufficient to induce epilepsy is supported by the fact that seizures were not reported in IGF-1 transgenic mice despite a 55% increase in brain size (62).

The FSDS phenotype that we describe is similar to two previously published conditions in FVB/N mice. Goelz *et al*. (63) describe spontaneous seizures in FVB/N mice that are accompanied by necrosis and astrocytosis. In contrast to the FSDS phenotype described here, the seizures occurred primarily in female mice, were frequently fatal and were not associated with an obvious enlargement of the brain. Hsiao *et al*. (64) also reported a spontaneously occurring atypical phenotype in FVB/N mice. These mice showed agitation, inactivity and seizures that were associated with astrocytosis in the hippocampus, amygdala, and cerebral cortex. Again, a dramatic enlargement of the brain was not reported.

Combined with our current findings this suggests that FVB/N mice may be particularly susceptible to the development of neurological abnormalities. In fact our previous work demonstrates that phenotypic severity in the YAC128 mouse model of HD is increased on the FVB/N background compared to C57BL/6 and 129 mice (65). Similarly, FVB/N mice have been shown to be more susceptible to excitotoxicity than other strains of mice (66). Thus, while using the FVB/N strain can be advantageous in exhibiting a more robust neurodegenerative phenotype, in future studies using this strain, it will be important to identify and exclude FSDS mice, especially in cases like YAC128 mice where the transgene of interest modulates the frequency of the FSDS phenotype.

### Huntingtin is beneficial in a neurological disorder independent of Huntington disease

We reported increased incidence of a FSDS phenotype with *Htt* +/− mice compared to their wild-type littermates, while the over-expression of wild-type HTT prevented the same phenotype. Our finding that huntingin protects against the development of a seizure disorder in FVB/N mice (**Fig. 7**) highlights the importance of wild-type huntingtin function in the brain and demonstrates that, in addition to moderating toxicity in HD (20, 21), huntingtin may be involved in compensatory mechanisms against neurodegeneration. (22) The importance of wild-type Htt for normal brain function is clearly supported by the fact that reducing Htt levels by at 50% or more results in neurological phenotypes (8, 13, 15, 16). Interestingly, Htt exhibits a significant increase in expression levels during development at postnatal day 7, which corresponds to a developmental decrease in sensitivity to excitotoxicity (67). There are reports of seizures in the neurodevelopmental disorder, LOMARS, a syndrome defined by putative *HTT* loss of function variants. (34) Epilepsy is also a common symptom of Wolf-Hirschhorn syndrome, a disease resulting from deletions in the short arm of chromosome 4 on which the *HTT* gene is located in humans (35). In addition, mouse models of the same disease, with deletions in the Htt gene, exhibit increased susceptibility to seizures (36).

**Figure 7.**
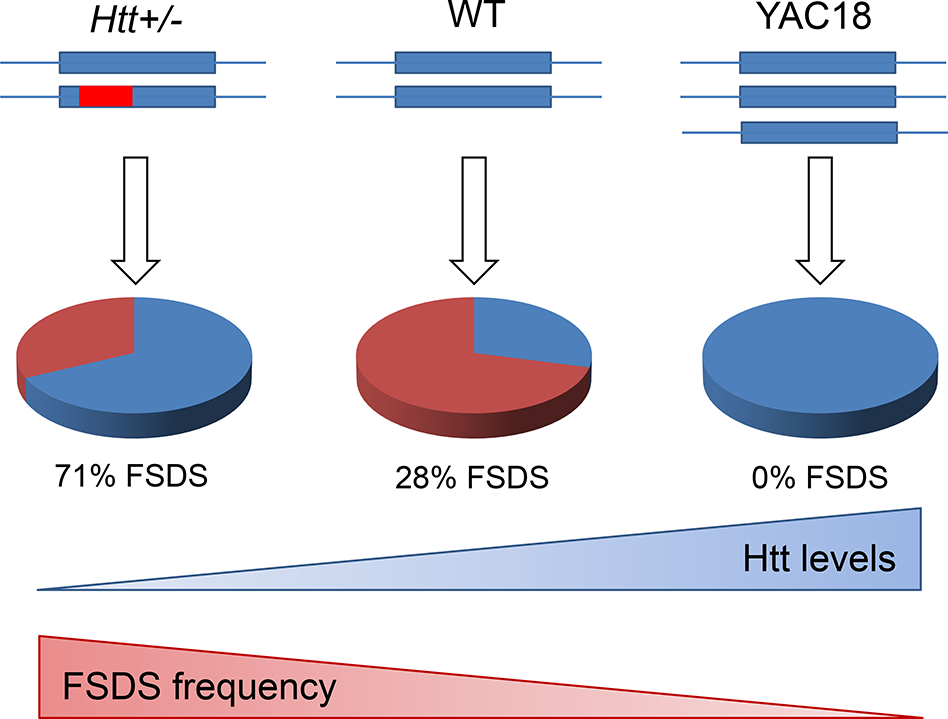
Huntingtin protects against epilepsy. A proportion of FVB/N mice develop an idiopathic seizure disorder characterized by megacephaly with features of epilepsy including poct-ictal behaviour, abnormal activity, spontaneous seizures, SUDEP (sudden unexpected death in epilepsy), neuronal hypertrophy, astrocytosis and upregulation of BDNF. We have named these mice FSDS mice (FVB/N Seizure Disorder with SUDEP). The frequency of the FSDS phenotype is modulated by the levels of full-length huntingtin (Htt). *Htt*+/− mice with 50% decreased levels of Htt have more than double the frequency of FSDS phenotype, while over-expression of HTT in YAC18 mice completely prevents the occurrence of the FSDS phenotype. Examination of the mechanism by which huntingtin protects against this epilepsy disorder reveals that huntingtin reduces seizure-induced damage but does not affect the number or severity of chemically induced seizures. These results suggest that reducing seizure-induced neuronal damage can limit the subsequent development of epilepsy.

### Polyglutamine expansion does not disrupt ability of huntingtin to protect against epilepsy

Previous work has demonstrated that polyglutamine expansion in the HTT protein disrupts only specific functions of HTT, but that other HTT functions remain largely unaffected. Mutant HTT can functionally replace wild-type Htt during development, as mice expressing only mutant human HTT with 128 glutamines exhibit normal development (20). Similarly, over-expression of both wild-type and mutant HTT have been shown to increase body weight (54). In contrast, Htt’s roles in promoting the expression, axonal transport, and uptake of BDNF are both disrupted by polyglutamine expansion (68–70). In this work, we show that the ability of huntingtin to protect against the FSDS epileptic phenotype is not disrupted by polyglutamine expansion as both wild-type and mutant HTT completely prevent the development of this disorder. Since mutant HTT fails to increase the expression or transport of BDNF, this suggests that the mechanism by which HTT prevents that development of the FSDS phenotype is not mediated by BDNF.

## Conclusions

In this work, we characterize a novel mouse model of epilepsy, which we call the <u>F</u>VB/N <u>S</u>eizure <u>D</u>isorder with <u>S</u>UDEP (FSDS) mouse model. Our results show that the levels of huntingtin expression modulate the frequency of the FSDS phenotype, indicating that huntingtin can protect against seizure disorders, at least partially due to its ability to reduce seizure-induced neurodegeneration. This work suggests the possibility that protective strategies aimed at increasing the levels of full-length huntingtin may be beneficial in some forms of epilepsy. In addition, our observation that decreasing wild-type huntingtin levels in the brain can have adverse effects has important implications for the development of huntingtin-lowering therapies for HD.

## MATERIALS AND METHODS

### Ethics Statement

All experiments were carried out in accordance with protocols approved by the UBC Committee on Animal Care and the Canadian Council on Animal Care.

### Mice

All mice were maintained on a pure FVB/N strain background (Charles River, Wilmington, MA; Jackson Laboratories, Bar Harbor, ME). YAC18 and YAC128 mice over-express wild-type or mutant human HTT, respectively, from a yeast artificial chromosome (YAC) (53, 55). *Htt*+/− mice are heterozygous for the targeted inactivation of the mouse *HTT* gene (*Htt*) (8). Shortstop mice express an N-terminal fragment of mutant HTT from a YAC transgene (56). Mice were group housed with a normal light-dark cycle in a clean facility and given free access to food and water. For all studies, experimenters were blinded to the genotypes of the mice.

### Identification of FSDS Mice

FSDS mice were identified using the following criteria: 1) observation of home cage immobility resembling post-ictal state of mice recovering from chemically-induced seizures, 2) observation of aggressive response to handling, 3) observation of megencephaly post-mortem. In cases where experiments were performed on the FSDS mice, the mice were identified by the first two criteria and the third criteria was examined after the completion of the experiment when the mice were killed. In every case when a mouse was identified as an FSDS mouse by the first two criteria, we observed megencephaly.

### *In vivo* electrophysiology

Two FSDS and three normal WT FVB/N mice of ages 7-9 months were used for EEG recording. The procedure consisted of three main steps: surgical implantation of electrodes, connection of the recording device, and data downloading and processing. All animal surgeries were carried out using aseptic techniques and in accordance with guidelines of the Canadian Council on Animal Care and approved protocol by the University of British Columbia Animal Care Committee.

Electrode implantation surgery

For electrode placement, animals were anesthetized with 3% isoflurane and placed in a stereotaxic frame. After exposing the cranium, four burr holes were drilled bilaterally over the frontal and parietal cortices (approximate bregma coordinates, frontal: AP = +1.5mm, ML = −/+1.8mm, parietal: AP = −2.4mm, ML = +/−2.2mm) and one over the occipital segment (approximate bregma coordinates AP = −5.03, ML = +0.6). Miniature stainless-steel screws (Part No. 0-80 X 1/16, Invivo1, USA) pre-soldered to insulted copper wire leads were screwed onto the skull holes with above coordinates to serve as EEG electrodes. Three electrodes (the two frontal and left parietal) were used to record EEG signals. The remaining two electrodes (right parietal and occipital) served as ground and reference electrodes, respectively. Wire terminals from these electrodes were connected to a 7-pin header that is mounted over the animal head. Screws and pin connector were further fixed in place by acrylic dental cement (Stoelting, USA). After surgery, mice were singly housed and allowed to recover for at least three weeks before proceeding with EEG recording.

### EEG recording

EEG recording was performed in freely-moving animals using Neurologger 2A (Evolocus, USA, http://www.evolocus.com/neurologger-2A.htm). This wireless non-telemetric system allows EEG data to be stored directly into a memory chip that is integrated within the head mount unit. It is also equipped with a 3-dimensional accelerometer that provides a simultaneous tracking of animal movement during EEG recording. To commence recording, Neurologger was connected to the implanted pin header with pre-set sampling rate of 400 Hz. EEG was continuously recorded in each animal for 24 - 43 hours.

### Data acquisition and analysis

At the end of each recording session, Neurologger was disconnected from the animal head and connected to a computer using Neurologger USB Adapter (Evolocus, USA). EEG and accelerometer data were downloaded offline from the logger memory to computer.

Retrieved data were then converted from binary to text or Float32IE formats. Data downloading and conversion were carried out using Downloader software tool version 1.27 (Evolocus, USA). Electrophysiological and accelerometer data were visualized and processed using EEGLAB versions 14.1.1 and 14.1.2 (71) running under Matlab versions R2017b/ R2019a (The MathWorks Inc., USA). Traces obtained from the three active EEG channels were plotted and analyzed in parallel with the synchronised animal acceleration data along the three orthogonal axes (x, y and z). EEG data were visually screened to identify potential convulsive seizure events. A convulsive seizure is defined by the presence of large-amplitude (> 2x baseline), high-frequency (> 5 Hz) discharges that last for minimum of 10 seconds and which is associated with sudden and vigorous changes in the animal movement along the three accelerometer axes.

### Behavioural Analysis

Open field activity was assessed in an automated open field apparatus (San Diego Instruments, San Diego, California). Activity was measured in the dark during the light cycle in a one hour open field trial. Activity was measured as the number of beam crosses in the trial. The open field chamber was cleaned with ethanol in between each open field trial. Beam crossing was used as an assessment of motor coordination and balance. Mice were placed at one end of a beam and the amount of time required for mouse to cross the beam to an escape chamber was measured. The maximum score on this test was 60 seconds. Mice falling off the beam were given a score of 60 seconds.

### Neuropathology

Mice were injected with heparin, terminally anesthetized by intraperitoneal injection of 2.5% avertin and perfused with 3% paraformaldehyde in phosphate buffered saline. Brains were post-fixed in 3% paraformaldehyde for 24 hours and then equilibrated with PBS. After weighing, brains were infiltrated with sucrose (25% in PBS) and frozen on dry ice before mounting with Tissue-TEK O.C.T. compound (Sakura, Torrance, CA). Twenty-five μm coronal sections were cut on a cryostat (Microm HM 500M, Richard-Allan Scientific, Kalamazoo, Michigan) and collected in PBS.

Brain cross-sectional area and neuronal size were determined with NeuN-stained coronal sections using Stereoinvestigator software (Microbrightfield, Williston, VA). Astrocytosis was assessed by glial fibrillary acidic protein (GFAP) staining using a Cy3-conjugated anti-GFAP antibody (1:200; C 9205, Sigma, Oakville, ON). Brain levels of Bdnf and Htt were determined by western blotting using protein lysates from either whole brain or microdissected brain sections with the following antibodies: Htt-specific MAB2166 antibody (1:2000, 1 hour, room temperature; Chemicon, Temecula, CA) or rabbit polyclonal anti-BDNF antibody (1:1000, overnight, 4 °C; Santa Cruz, Santa Cruz, CA).

### Chemically Induced Seizure Models

Pilocarpine (Sigma) and PTZ (Sigma) were used as chemical models of epilepsy. Pilocarpine injection at a concentration of 190 mg/kg (30 mg/ml solution), with a pre-injection of 0.1 mg/kg methylscopolamine (Sigma) induced SE in at least half of the injected mice with minimal mortality. Seven days after SE, the number of degenerating neurons in the hippocampus was determined by counting the total number of fluorojade-positive neurons. PTZ was administered 3 times per week for a total of 8 injections at a dose of 35 mg/kg. Following PTZ injection, mice were monitored for seizure activity until all of the injected mice had resumed normal home cage activity. Seizures were scored according to the following scale: 0 - no seizure, 1 - reduced activity, still, 2 - head nod, 3 - convulsive wave throughout body, 4 - full body seizure, rearing/in place, 5 - full body seizure, rolling, 6 - death.

### RNA Isolation and Quantitative RT-PCR

RNA samples from both FSDS (N=6) and normal FVB/N (N=3) mice were assessed for *Bdnf* variant expression levels using Quantitative RT-PCR (QRT-PCR). Total RNA was isolated from frozen mouse whole brains using the RNeasy Midi kit (Qiagen). QRT-PCR was performed according to a two-step process using the Quantitect Reverse Transcription Kit (Qiagen) for cDNA synthesis, and the Quantitect SYBR Green PCR kit (Qiagen). ABI Prism Software was used for real-time PCR analysis. Real-time PCR cycling conditions included a denaturation step of 95°C for 15 min, followed by 40 cycles of 95°C for 15 s, 60°C for 30 s, and 72°C for 30 sec. In both FSDS and normal FVB/N mice *Bdnf* variant 6 was not detectable by real-time PCR and not included in the results section.

### Chromatin Immunoprecipitation

Chromatin immunoprecipitation (ChIP) was performed following a modification of the Upstate Biotechnology ChIP kit protocol. Tissue was fixed in 1% formaldehyde. Cross-linked cell lysates were sheared by sonication in a 1% SDS lysis buffer to generate chromatin fragments with an average length of 100-200 bp. The chromatin was then immunoprecipitated using antibodies specific to acH3 which recognizes dimethylated Lys4 (Upstate Biotechnology, cat. # 07-030) or acH4 which recognizes acetylated Lys5, Lys8, Lys12 and Lys16 (Upstate Biotechnology, cat. #06-866) or an equivalent amount of control IgG (anti-rabbit, Santa Cruz, CA) at 4°C overnight. Protein-DNA-antibody complexes were precipitated with protein A-agarose beads coated with sheared salmon sperm DNA for one hour at 4°C, followed by two washes in low salt buffer, two washes in high salt buffer, two washes in LiCl buffer and three washes with 1x TE. The precipitated protein-DNA complexes were eluted from the antibody with 1% SDS and 0.1 M NaHCO_3_, then incubated at 65°C overnight in 200 mM NaCl to reverse formaldehyde cross-links. Following proteinase K digestion, phenol-chloroform extraction, and ethanol precipitation, samples were subjected to 40 cycles of PCR amplification using primer pairs specific for 200bp segments corresponding to the promoter region upstream of mouse *Bdnf* exon II.

Levels of specific histone modifications at the *BDNF* P2 promoter were determined by quantitative real-time PCR (StepOne Plus; Applied Biosystems, Foster City, CA). The following primers were used to amplify portions of the BDNF P2 promoter: 5’- GGATTTGTCCGAGGTGGTAG, −3’ and 5’- CAGCCTACACCGCTAGGAAG −3’. Input and immunoprecipitated DNA amplification reactions were run in triplicate in the presence of SYBR-Green (Applied Biosystems). Ct values from each sample were obtained using the StepOne v2 software. Relative quantification of template was performed as described previously (50) using the *ΔΔCt* method. Mean and SEM values were determined for each fold difference, and these values were used in two-tailed paired *t* tests to determine statistical significance (p < 0.05). Each PCR reaction, run in triplicate for each sample, was repeated at least two independent times.

### Spontaneous Seizures and Seizure Susceptibility

The frequency of seizures in the FSDS mice was determined by observing mice over a period of two hours. To assess the susceptibility of FSDS mice to audiogenic seizures, FSDS mice were exposed to 120 dB of noise for 1 minute. Mice were allowed to acclimatize to the noise chamber for 1 minute before the noise stimulation was applied. To assess the susceptibility of FSDS mice to pilocarpine-induced SE, mice were injected as described above.

### Seizure Induction

Mice were exposed to stressors including lipopolysaccharide (LPS), handling, and noise. For LPS treatment, FVB/N mice (N=12 control, N=17 LPS) were injected with 1 mg/kg LPS every 2 weeks from 6 to 9 months of age. Handling treatments consisted of repeat scruffing (15 sec.), novel cage (1 min.), and tail-suspension (15 sec.) 3 times a week, from 4-5 months of age (N=20 control, N=18 handled). Noise treatments included a 1-minute acclimation period before exposing mice to 120 dB of sound for 1 minute. Noise treatment was applied 3 times a week, for 1-month beginning at 21 days (N=11 control, N=20 noise) or 120 days (N=7 control, N=8 noise) of age. Following treatments FVB/N mice were monitored for the onset of the FSDS phenotype up to 12 months of age.

### Statistical Analysis

Longitudinal behaviour tests were analyzed using a repeated measures ANOVA. Comparisons between groups were performed using a 2-tailed student’s t-test. The Chi-square test was used to compare categorical data. Error is given as standard error of the mean. For all graphs * p<0.05, ** p < 0.01, *** p < 0.001.

## ACKNOWLEDGEMENTS

The authors would like to thank past and present members of the Leavitt Lab, and the Centre for Molecular Medicine and Therapeutics Transgenic facility (University of British Columbia, Vancouver, Canada) for their aid in this study.

JVR was supported by Canadian Institutes of Health Research (CIHR), Michael Smith Foundation for Health Research (MSFHR), Huntington Society of Canada, McGill Tomlinson Scholarships, Hereditary Disease Foundation, Parkinson’s Society of Canada, and Van Andel Research Institute. BRL was supported by CIHR, Huntington’s Society of Canada and the CHDI Foundation.

The authors declare no competing financial interests.

## MOVIE LEGENDS

**Movie 1.** Home cage behaviour of a normal FVB/N mouse. This movie illustrates the behavior of a normal FVB/N mouse. Notice that the mouse actively explores the cage.

**Movie 2.** Home cage behaviour of an FSDS mouse. This movie illustrates the behavior of an FSDS mouse. In contrast to a normal FVB/N mouse, these mice remain immobile for long periods of time and do not explore their surroundings.

**Movie 3.** Home cage behaviour of mouse following pilocarpine-induced *status epilepticus*. This movie illustrates the behavior of a mouse which has undergone status epilepticus following injection with pilocarpine. As with FSDS mice, these mice remain immobile for long periods of time and also do not explore their surroundings.

**Movie 4.** Spontaneous seizure observed in an FSDS mouse. This movie shows an FSDS mouse undergoing a spontaneous seizure. A normal FVB/N littermate walks past the seizing mouse.

**Movie 5.** Auditory SUDEP in an FSDS mouse. This movie shows an FSDS mouse undergoing SUDEP in response to an auditory stimulus. Before the auditory stimulus, the FSDS mouse is immobile. When the sound begins, the mouse responds by moving about the cage, first slowly then in a more frantic manner. Finally, the mouse exhibits a “popcorn” seizure and dies.

**Movie 6.** Comparison of normal FVB/N mouse and FSDS mouse in response to auditory stimulus. This movie compares the response of a normal FVB/N mouse and an FSDS mouse to an auditory stimulus. Before the sound begins, the normal FVB/N mouse is exploring the cage while the FSDS mouse is immobile. The auditory stimulus has no impact on the normal FVB/N mouse’s behavior. However, the auditory stimulus causes the FSDS mouse to move frantically about the cage, seize and then die.

